# Structural and dynamic insights into phosphate uptake by PHT1 transporter in rice

**DOI:** 10.1101/2025.08.04.668567

**Authors:** Zhangmeng Du, Zeyuan Guan, Hai Liu, Jie Zhang, Haitao He, Zhiwen Zheng, Wenhui Zhang, Lihuan Jiang, Jiaqi Zuo, Yan Liu, Beijing Wan, Haifu Tu, Faming Dong, Xuelei Lai, Lizhong Xiong, Ping Yin, Shaowu Xue, Yanke Chen, Zhu Liu

## Abstract

Phosphorus is an essential macronutrient for plants, primarily absorbed from the soil as inorganic phosphate (Pi) through root-located Pi transporters. Despite decades of research into these transporters as targets for developing Pi-efficient crops, their mechanisms for Pi import remain poorly understood. Here, we present the cryo-EM structures of the rice Pi importer OsPHT1;11 in both Pi-bound and unbound forms, characterize its conformational dynamics, and reveal how these dynamics contribute to its transport function. Pi is recognized through conserved residues found in plants, with its translocation facilitated by a typical alternating-access mechanism. SmFRET analyses reveal that this transporter undergoes dynamic conformational changes, which are differentially linked to its Pi transport capability, with a predominance of extracellular open conformations favoring Pi transport, while more populated intracellular open conformations hinder it. These findings provide insights into Pi uptake in plants and offer a foundation for designing genetically modified crops with improved phosphate efficiency.

## Introduction

Phosphorus (P) is one of the principal elemental nutrients for plants. A deficiency of this essential macronutrient severely limits plant growth, development, and crop productivity^1,2^. Plants utilize P primarily in the form of inorganic phosphate (Pi). Pi is absorbed by plant roots from the soil and is subsequently translocated to above-ground tissues, a process facilitated by Pi transporters located in the plasma membranes. These membrane proteins belong to the Phosphate Transporter 1 (PHT1) group, which is conserved across all major lineages of plants^3^. In rice, the PHT1 family consists of 13 members (OsPHT1;1-OsPHT1;13), with most highly expressed in roots under normal or Pi-deficient conditions^4–6^, and are transcriptionally regulated^7–9^. Notably, OsPHT1;11 (OsPT11) is specifically induced during arbuscular mycorrhizal (AM) symbiosis^5,10^, contributing to over 70% of Pi uptake in mycorrhizal rice^11^. This AM-specific Pi transporter is situated in the plasma membrane of cortical cells that envelop AM arbuscules, providing a site for symbiotic Pi uptake in rice and serving as an adaptive strategy to cope with limited availability of Pi in the soil^5,11,12^. Differential expression patterns and activity regulations of PHT1 members in various plant tissues highlight their specialized physiological roles in Pi uptake and tissue-to-tissue translocation^13,14^. Structurally, the PHT1 members of plants share remarkably similar amino acid sequences^15^ (Supplementary Fig. 1), indicating a conserved transport mechanism that facilitates the movement of Pi ions across the cell membrane.

Plant PHT1s belong to the phosphate: H^+^ symporters (PHS) within the major facilitator superfamily (MFS)^16^. MFS is the largest group of secondary active transporters that utilize electrochemical gradients to facilitate the movement of small molecules across biological membranes^17^. The canonical structure of MFS transporter comprises twelve transmembrane helices (TM1-TM12), which are divided into the N-domain (TM1-TM6) and the C-domain (TM7-TM12)^18^. These two domains organize a central cavity for substrate binding, positioned in the middle of the transporter across the biomembrane. MFS transporters employ a dynamic, alternating-access strategy to move substrates from one side of the membrane to the other^19^. This process involves the rearrangement of the N- and C-domains, transitioning between outward-open, occluded and inward-open conformational states^20,21^. The conformational transitions necessary for substrate transport are energetically driven by the movements of co-translocated ions down their concentration gradients^20,21^. In plants and fungi, phosphate transport is coupled with the co-translocation of H^+^ ions^22^. While the fundamental roles of plant Pi transporters have been extensively studied and explored as practical targets for developing Pi-efficient crop plants^14,23–25^, how these transporters recognize and transport Pi remain largely unknown. To date, the only experimental structure of an H^+^-coupled Pi transporter from the PHS family is that of the fungus *Piriformospora indica* (PiPT)^26^. The structure of Pi transporters in higher plant remains elusive, and the coordination of conformational dynamics that orchestrate the relative motions of the N- and C-domains is poorly understood. This limited mechanistic understanding hinders the rationale and efforts to develop Pi-efficient crops. Here we set out to determine the cryo-EM structure of OsPT11 from rice, visualize its conformational dynamics, and to establish how it recognizes and transports Pi across the cell membrane.

## Results

### Function characterization and structure determination of OsPT11

Plants employ two pathways for Pi uptake: direct acquisition from the soil via roots and indirect translocation from symbiotic fungi. High-affinity Pi transporters, such as the low-Pi induced OsPT6 and OsPT8^27,28^, are responsible for direct Pi uptake under Pi-deficient conditions in the soil^15,29^. However, the Pi transport capability of the AM-specific OsPT11 for indirect symbiotic Pi uptake is not clear. To assess the function of OsPT11, we performed Pi transport assays in a heterologous yeast system. We utilized a yeast mutant strain (YP100) that lacks Pi transporters and cannot grow on Yeast Nitrogen Base (YNB) medium^30^. As a proof-of-concept, we complemented this strain with two yeast Pi transporter genes separately, the high-affinity transporter *PHO84* and low-affinity transporter *PHO91*, and assessed yeast growth on media containing different Pi concentrations. Consistent with their Pi transport capabilities, transformation of the yeast with *PHO84* restored growth effectively, while *PHO91* had a lesser effect on growth complementation (Fig. 1A,B). In parallel, we observed that *OsPT11* partially complemented yeast growth, with a slower rate than *PHO91* (Fig. 1A,B). These results suggest that OsPT11 transport Pi into yeast but likely functions as a low-affinity Pi transporter. Low-affinity Pi transporters induced by AM fungi have also been identified in plant crops^31–33^.

**Fig. 1.**
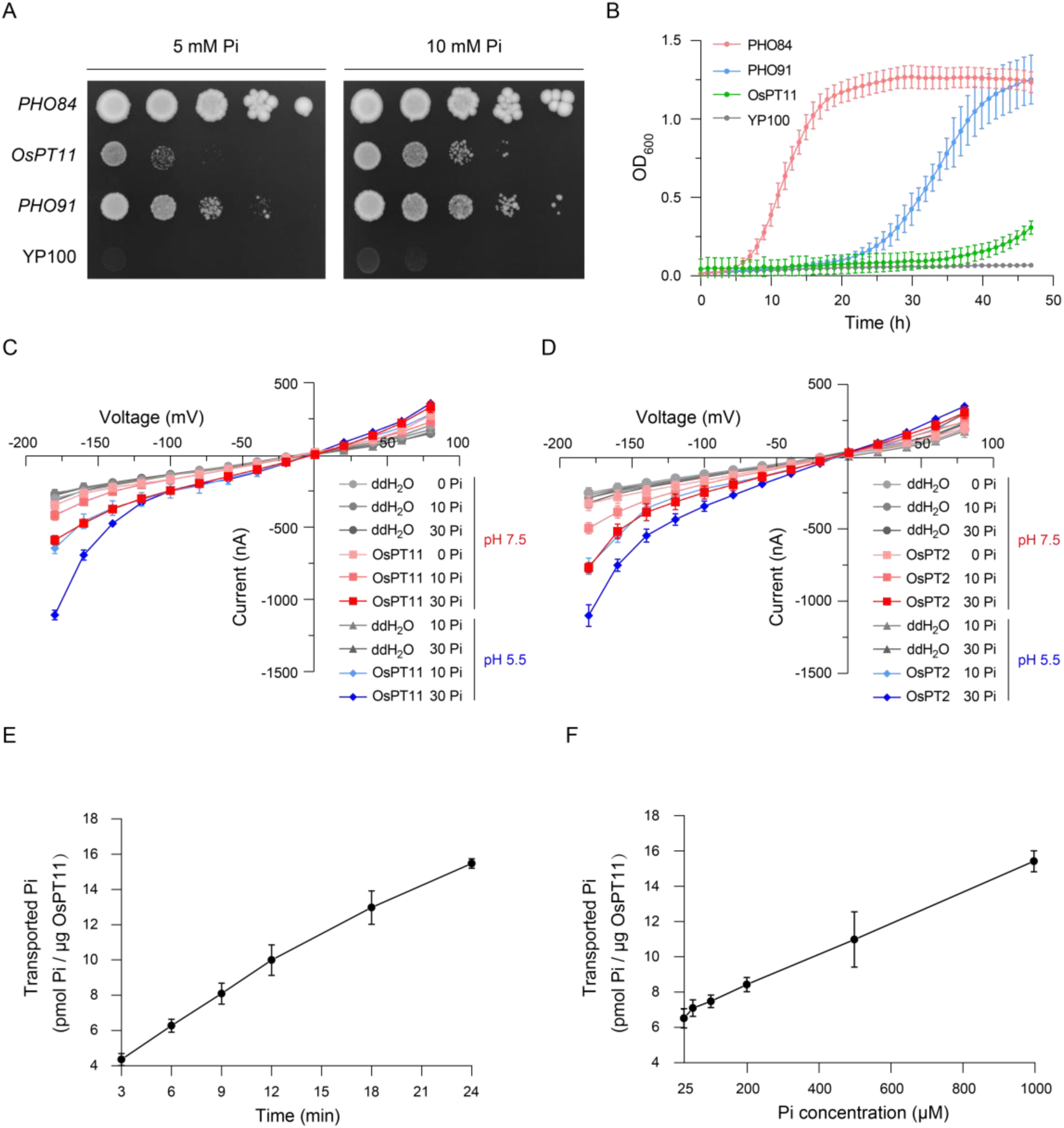
Function characterizations. **A-B** Functional complementation analysis of the Pi uptake-defective yeast mutant YP100 expressing various Pi transporters, assessed in solid YNB medium (**A**) and liquid medium (**B**), respectively. Yeast growth curves are represented as mean ± SD from 5 independent experiments, cultured in the presence of 10 mM phosphate. **C-D** Electrophysiological properties of OsPT11 (**C**) and OsPT2 (**D**), respectively. The current-voltage relationships of the TEVC recordings are shown. The voltage steps ranged from 80 to -180 mV in 20 mV decrements. The current was recorded in *Xenopus* oocytes injected with water in 0 mM Pi (n=4, pH 7.5), 10 mM Pi (n=4, pH 7.5; n=4, pH 5.5), 30 mM Pi (n=4, pH 7.5; n=4, pH 5.5), expressing OsPT11 in 0 mM Pi (n=8, pH 7.5), 10 mM Pi (n=6, pH 7.5; n=7, pH 5.5), 30 mM Pi (n=8, pH 7.5; n=10, pH 5.5), and expressing OsPT2 in 0 mM Pi (n=8, pH 7.5), 10 mM Pi (n=9, pH 7.5; n=7, pH 5.5), 30 mM Pi (n=8, pH 7.5; n=10, pH 5.5). Data shown are means ± SEM. **E-F** The accumulation of transported Pi in proteoliposomes carrying OsPT11 is related to the duration of the transport reaction (**E**) and the external Pi concentration (**F**), respectively. The time-course activity was collected at an external Pi concentration of 50 μM, with transport reaction times of 3, 6, 9, 12, 18, and 24 minutes. The Pi concentration-dependent activity was measured with a 6-minute reaction duration, using external Pi concentrations of 25, 50, 100, 200, 500, and 1000 μM. These transport activity data are represented as mean ± SEM from 3-5 independent experiments.

To further investigate the properties of OsPT11, we conducted electrophysiological experiments using *Xenopus laevis* oocytes. When Pi was added to the bath solution, oocytes injected with OsPT11 cRNA exhibited a significant increase in voltage-dependent membrane currents compared to the negative control oocytes injected with water (Fig. 1C). These observations indicate that OsPT11 facilitates the transport of Pi across the oocyte membrane. Additionally, as OsPT11 is an H^+^-coupled symporter, acidification of the bath solution resulted in an increase in Pi-induced currents (Fig. 1C). The currents observed in oocytes expressing OsPT11 were dependent on the concentration of Pi in the bath solution, requiring tens of millimolar Pi (Fig. 1C). These electrophysiological properties align with those of OsPT2 (Fig. 1D), a reported low-affinity Pi transporter^27^. These results suggest that OsPT11 most likely acts as a low affinity Pi transporter, corroborating our findings from the heterologous yeast assay (Fig. 1A,B).

We then biochemically characterized the Pi transport capability of OsPT11 in vitro. By transiently expressing OsPT11 in HEK293F cells, we purified the protein and obtained the detergent-solubilized sample with good behavior (Supplementary Fig. 2 and Methods). We reconstituted this sample into liposomes and conducted Pi transport assays (Methods). The results showed that OsPT11 exhibited Pi transport activity, facilitating the transport of Pi into the proteoliposomes in a manner dependent on external Pi concentration and the duration of the transport reaction (Fig. 1E,F). The Pi accumulation in the proteoliposomes showed a nearly linear correlation with the external Pi concentration up to 1 mM in the established assays (Fig. 1F). This implies that OsPT11 may operate as a low-affinity Pi transporter with a *K*_m_ in the millimolar range, consistent with our findings from the heterologous assays conducted in yeast and *X. laevis* oocyte systems. This low-affinity transport property of OsPT11 aligns with its physiological role in arbuscular mycorrhizal rice, where it facilitates the acquisition of Pi released by the associated fungi^5,10,11^.

To resolve the structural basis of OsPT11 for Pi recognition and transport, we sought to perform Cryo-EM analysis. OsPT11 was prepared in digitonin detergent in the presence of 25 mM phosphate at two pH conditions, pH 5 and pH 8, and high-quality cryo-EM images were collected for single-particle reconstruction (Methods). The 3D reconstructed density maps were refined to 3.4 Å for OsPT11 prepared under both conditions (Fig. 2A, Supplementary Fig, 3). The well-resolved EM density allowed us to build the atomic models, which we referred to as OsPT11^pH5^ and OsPT11^pH8^ (Supplementary Fig. 4, Supplementary table 1). The transporter structure exhibits a dimeric conformation with minimal contacts between the two protomers (Fig. 2A, Supplementary Fig. 4A), suggesting that they may function as independent units. The protomers of OsPT11^pH5^ and OsPT11^pH8^ adopt a similar overall fold, with a RMSD of 0.57-0.66 Å. Closer inspection of the structural density reveals a putative phosphate ion bound to one protomer of OsPT11^pH5^ (Supplementary Fig. 4C), while it is not visible in the other protomers of OsPT11^pH5^ and OsPT11^pH8^. This observation is reminiscent of our function characterizations that OsPT11 operates as a low-affinity Pi transporter, which may explain the rarity of Pi capture in the Cryo-EM structures, despite the sample was prepared in the presence of 25 mM phosphate. In this manuscript, we use the structures of the Pi-bound protomer of OsPT11^pH5^ and the Pi-free protomer of OsPT11^pH8^ for further mechanistic analyses.

**Fig. 2.**
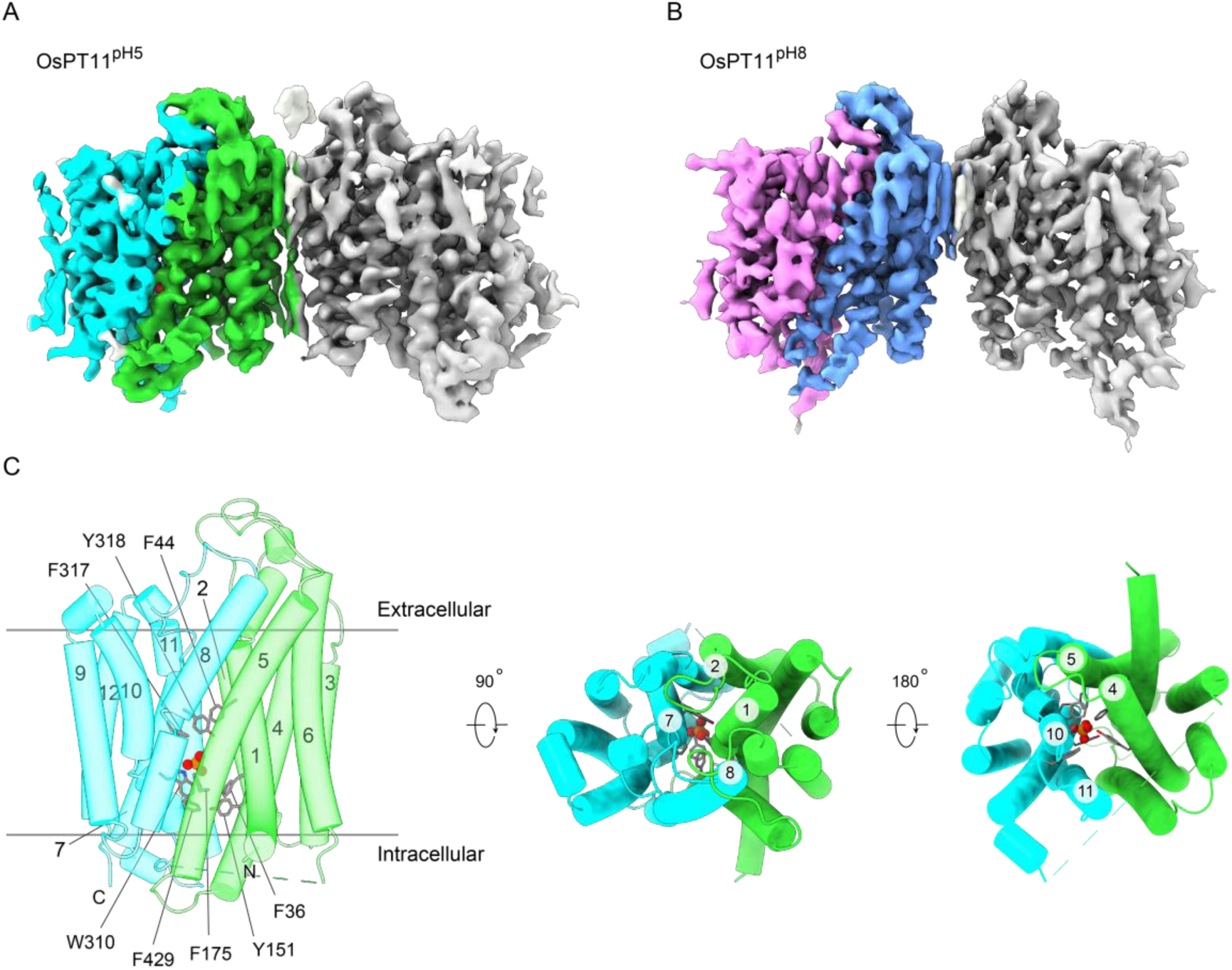
Cryo-EM structures of OsPT11. **A-B** The EM density maps of OsPT11 determined at pH 5 (**A**) and pH 8 (**B**) conditions, respectively. OsPT11 forms dimers in the micelles at the two conditions, with lipid-like densities are observed between protomers. **C** The structure of Pi-bound OsPT11. The N-domain (TM1-TM6) and C-domain (TM7-TM12) are colored in green and cyan, respectively. The phosphate ion is represented as a sphere. Residues F44, F317, and Y318 block the entry pathway to the Pi binding site, while residues F36, Y151, F175, F429, and W310 obstruct the cytosolic exit of Pi, represented as sticks.

### Structural basis for phosphate recognition and transport

The structure of Pi-bound OsPT11 adopts a canonical MFS transporter fold, with the N-domain (TM1-TM6) and C-domain (TM7-TM12) associated and positioned perpendicular to the membrane (Fig. 2C). In this configuration, the Pi ion is coordinated within a central cavity at the interface between the N- and C-domains. The extracellular side features interactions between TM1-2 and TM7-8, blocking the entry pathway to the Pi binding site. Concurrently, TM4-5 are positioned near TM10-11 on the intracellular side, obstructing the cytosolic exit of Pi. Specifically, aromatic residues F44 (in TM1), F317 and Y318 (in TM7) form a hydrophobic cluster above the Pi binding site, occluding the entry pathway. Meanwhile, other aromatic residues, including F36 (in TM1), Y151 (in TM4), F175 (in TM5), W310 (in TM7), and F429 (in TM10), are positioned below the binding site, obstructing the exit pathway for Pi ion. These residues are fully conserved among plant PHT1 members, indicating their critical gating roles (Supplementary Fig. 1). As the centrally bound phosphate remains inaccessible from both the extracellular and intracellular sides, this Pi-bound OsPT11 structure represents an occluded intermediate within the framework of the alternating-access transport model.

The Pi-bound OsPT11 structure provides a molecular basis for understanding how OsPT11 recognizes and transports phosphate. Within the central cavity, the Pi ion is coordinated by a series of specific residues through polar interactions (Fig. 3A, Supplementary Fig. 4C). The side chains of Q178, W310, D314, and N425 form hydrogen bonds with the Pi, while K453 contributes to Pi binding through electrostatic interactions. All these Pi-recognizing residues are 100% conserved in plant PHT1 transporters (Supplementary Fig. 1). This configuration of the Pi binding site in OsPT11 is similar to that in fungus PiPT^26^, except that the homologous residue K459 in PiPT, which corresponds to K453 in OsPT11, does not participate in Pi binding (Supplementary Fig. 5). Substitutions of side chains in Pi-recognizing residues, including Q178, W310, D314, N425, and K453, with alanine resulted in impaired Pi transport activity (Fig. 3B), highlighting crucial roles of these residues in Pi transport.

**Fig. 3.**
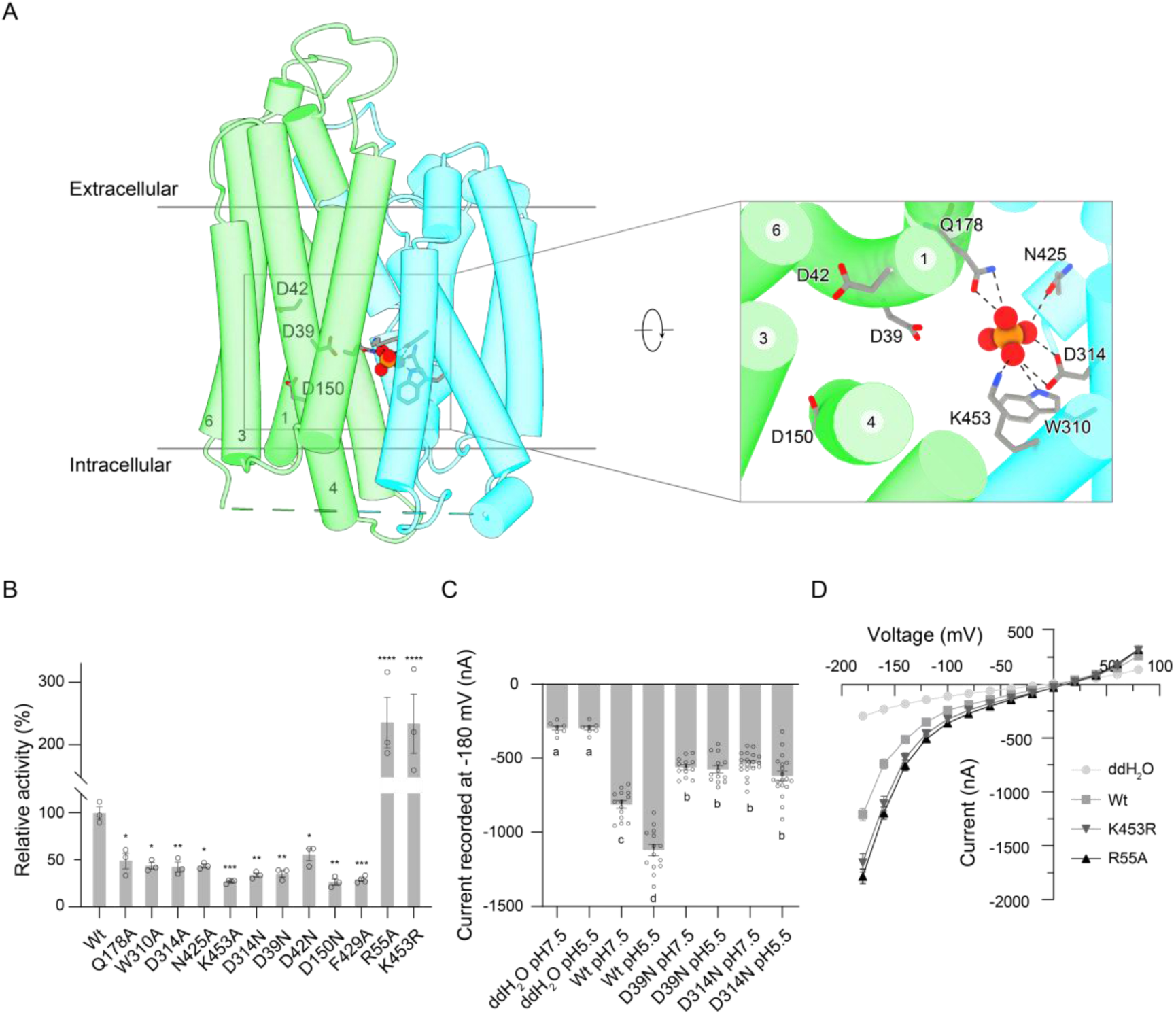
Structural basis of OsPT11 for phosphate recognition and transport. **A** Pi recognition site. The black dashed lines indicate plausible interactions between the Pi ion and the protein. Acidic residues D39, D42 and D150 lined near the Pi-binding site are represented as sticks. **B** Proteoliposome-based Pi transport assays. These experiments were conducted at the external Pi concentration of 50 μM, with a transport reaction time of 6 minutes. The transport activity is determined by measuring the accumulation of phosphate in the proteolipsoomes. Relative transport activity of each OsPT11 mutant is referenced to the wild-type. Data are represented as mean ± SEM from 3-4 independent experiments. Statistical analysis: One-way ANOVA. \**p* < 0.1, \*\**p* < 0.01, \*\*\**p* < 0.001, \*\*\*\**p* < 0.0001. **C-D** Electrophysiological properties of OsPT11 and variants. Currents (**C**) were recorded at −180 mV in *Xenopus* oocytes injected with water (n=7, pH 7.5; n=7, pH 5.5), expressing OsPT11 (Wt, n=14, pH 7.5; n=14, pH 5.5), D39N mutant (n=13, pH 7.5; n=13, pH 5.5), D314N mutant (n=19, pH 7.5; n=19, pH 5.5) in the bath solution with 30 mM Pi. Data are represented as mean ± SEM. Different letters represent *p* < 0.05; One-way ANOVA. Current-voltage relationship (**D**) was recorded in oocytes injected water (n=6), expressing OsPT11 (Wt, n=13), R55A mutant (n=31), K453R mutant (n=21) in the presence of 30 mM Pi at pH 5.5. Data are represented as mean ± SEM.

In addition to the basic residue K453 in the Pi-binding site of OsPT11, it is interesting that an acidic residue (D314) is also present, which likely contributes to binding the anionic Pi ion. Acidic residues in the Pi-binding site have also been observed in the channel-like Pi exporter XPR1 in humans^34–36^, the Na^+^-coupled Pi importer from *Thermotoga maritima*^37^, and the fungal H^+^-coupled Pi-importer (PiPT)^26^. Previous molecular dynamics simulations of PiPT suggested that protonation of D324 (corresponding to D314 in OsPT11) is required for Pi binding, while the deprotonation triggers the release of the anionic substrate. Consistent with this conserved implication, we observed that Pi bound to OsPT11 at low pH (pH 5) but not at high pH (pH 8) in our cryo-EM structures. The acidic environment might protonate D314 in OsPT11, promoting Pi binding, while an alkaline pH led to deprotonation and thus a Pi-free (*apo*) state. Therefore, the reversible protonation-deprotonation of D314 is critical for the Pi transport cycle. Supporting this, alanine substitution of D314 or a protonation mimic (D314N) largely reduced the transport activity of OsPT11 (Fig. 3B,C). D314 is conserved among plant PHT1 transporters (Supplementary Fig. 1), indicating functional and mechanistic conservations. This is further evidenced by the reduced Pi-induced currents observed in the D305N mutation of OsPT2 (corresponding to D314 in OsPT11) in electrophysiological experiments (Supplementary Fig. 6A).

OsPT11 is a phosphate:H⁺ symporter that harnesses energy from H⁺ co-translocation to transport phosphate. Proton translocation in transporters is typically facilitated by a series of negatively charged residues. In OsPT11, three conserved acidic residues— D39, D42, and D150 (Supplementary Fig. 1)—are lined near the Pi-binding site, forming a tunnel surrounded by TM1, TM3, TM4, and TM6, which extends toward the cytosol (Fig. 3A). This suggests their involvement in H^+^-energized Pi transport. Proteoliposome-based transport assays showed that mutations D39N, D42N or D150N impaired Pi transport activity (Fig. 3B). Notably, D39 is closest to the Pi-binding site, likely serving as a relay for proton transfer between this site and the other acidic residues (Fig. 3A). Electrophysiological experiments further confirmed that the D39N mutation significantly reduced Pi influx, irrespective of whether the bath pH was acidic or basic (Fig. 3C). This feature is conserved, as evidenced by the electrophysiological properties of the homolog OsPT2 with a D35N mutation (corresponding to D39 in OsPT11) (Supplementary Fig. 6A).

The overall structure of *apo* OsPT11 shows little difference compared to the Pi-bound occluded structure (Supplementary 7), with the central cavity being inaccessible from both membrane sides. We propose that the *apo* form represents an empty occluded conformational state, a typical intermediate in the transport cycle through which a MFS transporter rests itself to outward-facing conformation after substrate release^21^. These structural findings indicate that OsPT11 employs similar conformational states to facilitate both outward-to-inward and inward-to-outward transitions during the transport cycle. In the central cavity, the side chain of K453 points away from the Pi binding site in the *apo* form, but interacts with the Pi ion in the Pi-bound state (Supplementary Fig. 7). This alteration in K453 side chain orientation suggests its crucial role in recognizing and transporting the substrate. We hypothesized that replacing K453 with a more alkaline residue, such as arginine, which possesses a higher p*K*_a_, would enhance Pi binding and consequently increase transport activity. To validate this hypothesis, we conducted transport assays and electrophysiological experiments to assess the Pi transport activity of OsPT11 carrying the K453R substitution. The results showed that the K453R substitution significantly enhanced Pi transport in OsPT11 (Fig 3B,D). Additionally, a similar enhancement was observed in the homolog OsPT2 with the K449R substitution (corresponding to K453 in OsPT11) (Supplementary Fig. 6B), further supporting our hypothesis regarding the role of this basic residue in phosphate transport.

### Dynamics of conformational transitions

To achieve alternating-access transport, a transporter transitions between outward-open and inward-open conformations, with the N- and C-domains alternatively opening at the extracellular and intracellular sides. To investigate the conformational dynamics of OsPT11 and their role in phosphate transport, we performed single-molecule fluorescence resonance energy transfer (smFRET) analysis, a method to characterize protein dynamics at single-molecule level in solution^38–40^. To monitor the relative motion of the N- and C-domains, we introduced FRET pairs on the extracellular and intracellular sides of OsPT11, respectively. In a cysteine-free OsPT11 background (Methods), the extracellular FRET pair was created by labeling fluorophores at K130C (in the N-domain) and T404C (in the C-domain) (Fig. 4A), while the intracellular pair was labeled at K242C (in the N-domain) and A296C (in the C-domain) (Fig. 4D). Our occluded cryo-EM structure shows that contacts between TM1-TM2 and TM7-TM8 close the N- and C-domains at the extracellular side (Fig. 2C, Fig. 4A), while contacts between TM4-TM5 and TM10-TM11 close the domains at the intracellular side (Fig. 2C, Fig. 4D). Based on this, we hypothesized that a low FRET efficiency would occur if the N- and C-domains move apart, opening the transporter. The smFRET results revealed that both FRET pairs detected dynamic conformational transitions of OsPT11 (Fig. 4B,E). The FRET efficiency fluctuations and their distributions suggest that OsPT11 primarily samples two inter-converting conformational states (Fig. 4B,E). The low-FRET species represent the open state of the N- and C-domains, owing to an increased distance between the fluorophores. In contrast, the high-FRET species correspond to the closed state, where the domains are in proximity. On the extracellular side, the high-FRET species had an efficiency centered around 0.62 (Fig. 4B), corresponding to a distance of 57.1 Å between the FRET pair (*R*_0_ = 62 Å^41^). This distance is aligned with the theoretical value of 60.1 ± 5.2 Å calculated from our occluded cryo-EM structure (Supplementary Fig. 8A), corroborating the structural and smFRET findings. Similarly, the high-FRET species on the intracellular side, centered at ∼0.66 (Fig. 4E), corresponded to a distance of 55.5 Å, which is also aligned well with the simulated value of 53.5 ± 5.9 Å derived from the experimental structure (Supplementary Fig. 8B). Together, these results demonstrate that the conformations of OsPT11 are dynamic in solution, with the N- and C-domains transitioning between open and closed states on both sides of the transporter, a hallmark feature essential for its transport function.

**Fig. 4.**
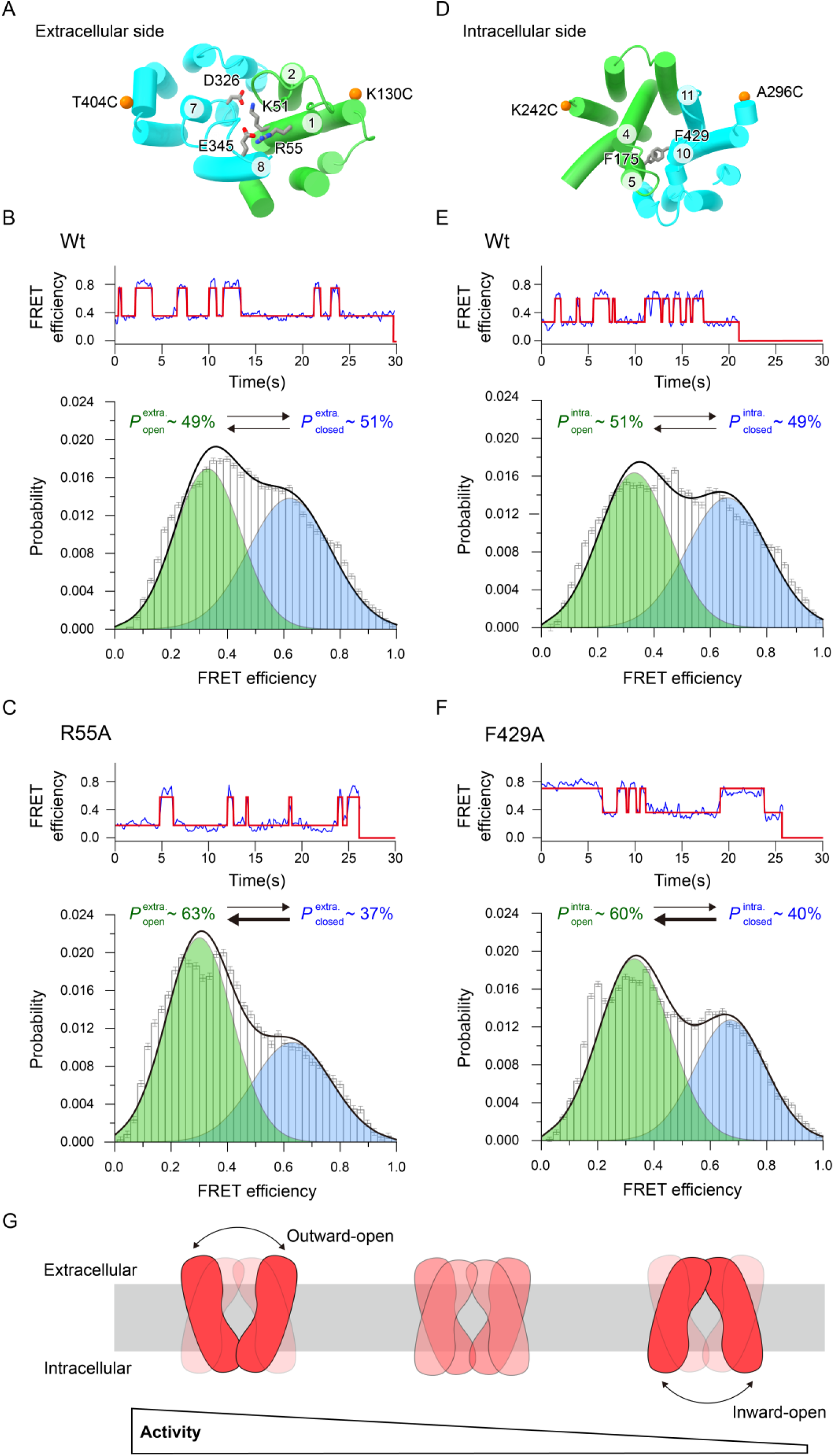
Structural dynamics of OsPT11. **A** Extracellular cut-view of the occlude structure. The orange spheres indicate the sites for fluorophores conjugation. **B** SmFRET efficiency distribution of OsPT11 observed from the extracellular FRET pair. Top: An exemplary smFRET trace and analysis using a two-state hidden Markov model. Bottom: The smFRET efficiency distribution (gray histogram) can be fitted to two Gaussian peaks, representing two FRET species. The Low-FRET species (colored green) and high-FRET species (colored blue) correspond to the extracellular open and closed states, with a population of 49% and 51%, respectively. The black line represents the cumulative fitted distributions. The length of the error bars represent the SD of data in each bin. **C** SmFRET efficiency distribution of the R55A OsPT11 mutant observed from the extracellular FRET pair. **D** Intracellular cut-view of the occlude structure. The orange spheres indicate the sites for fluorophores conjugation. **E** SmFRET efficiency distribution of OsPT11 observed from the intracellular FRET pair. **F** SmFRET efficiency distribution of the F429A OsPT11 mutant observed from the intracellular FRET pair. **G** A model of the OsPT11 conformational dynamics differentially linked to its transport function. A predominance of extracellular open conformations favors Pi transport, whereas a greater population of intracellular open conformations hinder it.

We next investigated how the conformational dynamics of OsPT11 is relate to its transport function. In the occluded intermediate, residues K51 and R55 in TM1 likely interact with residues E345 and D326 in TM7-TM8, which close the extracellular side (Fig. 4A). We hypothesized that disrupting these interactions would open the transporter on this side. To test this, we introduced the R55A substitution and performed smFRET analysis using the extracellular FRET pair. The results showed that the R55A substitution shifted the conformations equilibrium toward the low-FRET species, increasing its population to ∼63% (from ∼49%), while decreasing the high-FRET species (Fig. 4B,C). This suggests that the R55A substitution attenuates the TMs interaction, favoring a greater proportion of extracellular open conformations. To understand how this conformational redistribution affects transport function, we measured the Pi transport activity of the R55A mutant. Both proteoliposome-based assays and electrophysiological experiments revealed a significant increase in Pi transport (Fig. 3B,D), suggesting that a higher proportion of outward-open conformation enhances transport activity. Given that alternating-access transport relies on negative cooperativity between the outward- and inward-open states, preventing their simultaneous opening^21,42^, we speculated that, in contrast to the effects of the predominance of outward-open conformation, an increased proportion of the inward-open conformation would impair Pi transport. To test this hypothesis, we introduced the F429A mutation in TM10, which is expected to disrupt a π-π interaction with F175 in TM5 on the intracellular side (Fig. 4D), potentially destabilizing the closed intermediate. SmFRET analysis of the intracellular FRET pair showed that the F429A mutation enriched the low-FRET species (Fig. 4E,F), indicating a preference for the intracellular open conformation. This conformational shift of F429A mutant was in line with the expected outcome, corresponding to a decrease in Pi transport activity (Fig. 3B,D). Together, these findings demonstrate that the conformational dynamics of OsPT11 are closely linked to its transport function (Fig. 4G), with a predominance of extracellular open conformations favoring Pi transport, whereas more populated intracellular open conformations hindering Pi transport.

## Discussion

As an essential macronutrient and indispensable metabolite for plants, phosphate’s absorption from the soil, tissue-to-tissue translocation, and homeostasis maintenance within cells are fundamental to plant growth and development. These processes are facilitated by a range of Pi transporters, including the plasma membrane-localized PHT1 and PHO1 families^6,43–46^, as well as intracellular transporter families such as PHT2, PHT3, PHT4, PHT5, and SPX-MFS^46–51^. In this study, the experimental structure and dynamics characterization of OsPT11 (OsPHT1;11) provides the first snapshots for understanding phosphate recognition and transport by a rice Pi transporter. Since the discovery of the first plant phosphate transporters^43^, decades of extensive efforts have been made to develop genetically engineered crops with improved phosphate efficiency^1,2,46,52,53^, through the overexpression of specific Pi transporter genes. Here, our mechanistic studies on Pi transport demonstrate that its activity is linked to intrinsic dynamics; moreover, this activity can be modulated by redistributing the conformational space. These findings may offer new strategies for modifying Pi transporters by fine-tuning their functions, rather than simply relying on genetic overexpression.

Plant Pi transporters exhibit functional redundancy and vary in their transport capabilities to adapt to fluctuations in Pi availability in the environment. These transporters are characterized as either low- or high-affinity Pi transporters, depending on their transport capacity. For example, the rice PHT1 family consists of 13 members, while *Arabidopsis* contains 9 PHT1 transporters. An interesting open question is what determines their differing Pi transport capabilities, given that PHT1s share a conserved amino acid sequence, particularly in the residues of the Pi-binding site (Supplementary Fig. 1). Our studies show that the Pi transport activity of OsPT11, a low-affinity transporter, is higher when the outward-open conformation is more populated in solution and lower when the inward-open conformation predominates. This suggests that transport activity is tightly linked to the distribution of specific conformational states within the alternative-accessing MFS transporter. Whether other low- and high-affinity transporters harbor distinct conformational distributions, and if these intrinsic dynamics contribute to their differences in activity, remains an important and intriguing question for future study.

## Methods

### Protein expression and purification

Rice OsPHT1;11 (OsPT11) DNA was amplified from the rice cDNA library, and subcloned into a pMlink vector containing an N-terminal 3×Flag tandem affinity tag. The site-specific mutations were introduced by overlapping PCR and were verified by DNA sequencing.

Proteins were expressed in Expi293F mammalian cells (Invitrogen) via transient transfection. Cells were cultured in Union-293 media (Union-Biotech, Shanghai) and were transfected at a density of 2.0×10^6^ cells per milliliter. For a 1-liter cell culture, 2 mg of plasmids were premixed with 4 mg of 4 kDa linear polyethylenimine (PEI) (Polysciences) in 50 ml of medium and incubated for 20 minutes. This mixture was then added to the cell culture. Transfected cells were cultured at 37 °C under 5% CO_2_ for an additional 60 hours before harvesting.

For the preparation of the cryo-EM sample, transfected cells were collected by centrifugation at 800 g for 20 minutes. The pellets were resuspended in TBS buffer containing 50 mM Tris-HCl (pH 7.4), 150 mM NaCl, 1% (w/v) LMNG (Anatrace), 0.1% (w/v) CHS (Anatrace), and 0.25% (w/v) Soy Phospholipids (Sigma). After incubation at 4 °C for 1.5 hours, the supernatant was centrifuged at 23,000 g for 40 minutes and then incubated with anti-Flag G1 affinity resin (Genscript) at 4 °C for 40 minutes. The resin was rinsed with 30 bed volumes of wash buffer containing 50 mM Tris-HCl (pH 7.4), 150 mM NaCl, and 0.06% (w/v) digitonin (Anatrace). Proteins were eluted using wash buffer supplemented with 250 μg/ml Flag peptide (Genscript). The eluent was concentrated and applied to size-exclusion chromatography (SEC) using a Superose-6 Increase 10/300 column (GE Healthcare) in a buffer containing 25 mM phosphate (pH 8.0), 150 mM NaCl, 1 mM EDTA, 5 mM DTT, and 0.06% (w/v) digitonin (Anatrace). For the determination of OsPT11^pH5^ structure, the sample was prepared in a buffer containing 25 mM phosphate (pH 5.0), 150 mM NaCl, 1 mM EDTA, 5 mM DTT, and 0.06% (w/v) digitonin (Anatrace). The peak fractions of OsPHT11 were collected and pooled to approximately 6.5 mg/ml for cryo-EM grids preparation

Samples used for the proteoliposome-based transport assays were extracted from transfected cells using a buffer containing 50 mM Tris-HCl (pH 7.4), 150 mM NaCl, and 1% (w/v) n-Dodecyl-β-D-Maltopyranoside (DDM). The targets were further purified using anti-Flag G1 affinity resin, followed by a Superose-6 Increase 10/300 column. The purified proteins were then prepared in a buffer containing 25 mM Tris-HCl (pH 8.0), 150 mM NaCl, and 0.02% (w/v) DDM for proteoliposome reconstitution.

To prevent intramolecular FRET due to the dimerization property of OsPT11, a co-expression strategy was employed for fluorescent dye conjugation and intermolecular smFRET analysis. Two OsPT11 plasmids were used: one containing an N-terminal 3×Flag tag (labeling plasmid) and the other containing an additional Avi-tag ahead of the N-terminal 3×Flag tag (immobilizing plasmid). In both plasmids, the four intrinsic cysteines (C45S/C117M/C137M/C444S) were substituted to generate a cysteine-free background. Two additional cysteines, K130C/T404C (or K242C/A296C), were introduced in the labeling plasmid for specific dye labeling. After co-expression, the protein was purified following the same protocol as for the cryo-EM sample, with the addition of 2 μM BirA, 5 mM MgCl_2_, 4 mM ATP, 1 mM biotin, and 2 mM DTT during incubation with anti-Flag G1 affinity resin (Genscript) for biotinylation. The incubation was carried out for 2 hours. The protein was then purified and prepared in a buffer containing 25 mM Tris-HCl (pH 8.0), 150 mM NaCl, and 0.06% (w/v) digitonin for subsequent use.

### Cryo-EM grid preparation and data acquisition

3.5 μl aliquots of the purified OsPT11 protein was dispensed onto glow discharged holey carbon grid (Quantifoil Cu R1.2/1.3, 300 mesh). The grid was blotted with a Vitrobot Mark IV (ThemoFisher Scientific) using 3.5 s blotting time with 100% humidity and 8 °C, and was plunge-frozen in liquid ethane cooled by liquid nitrogen. The cryo-grids of OsPT11^pH8^ were transferred to 300 kV Titan Krios electron microscopes (Thermo Fisher), equipped with a GIF Quantum energy filter (slit width 20 eV) and a Gatan K2 Summit detector. Micrographs were recorded in the super-resolution mode with a magnification of 130,000×, corresponding to a pixel size of 1.08 Å. For the data collection of OsPT11^pH5^, movies were collected on a 300Kv Titan Krios electron microscopes, equipped with a Bioquantum energy filter and a Gatan K3 direct electron detection. Dose-fractionated images were recorded at a magnification of 81,000×, with a super-resolution pixel size of 1.07 Å. All datasets were acquired using EPU software, with a defocus range set between −1.2 and −2.2 µm. The total dose rate on the detector was 50 e^−^/Å^2^, and each micrograph stack contained 32 frames.

### Cryo-EM data processing

A diagram of the procedures for data processing are described in Supplementary Fig. 3. For the OsPT11^pH5^ analysis process, a total of 3,017 movies were collected. After motion correction and CTF estimation^54,55^, all micrographs were subjected to subsequent processing. The results of two-dimensional (2D) classification indicated that the particle features exhibited dimer characteristics. After multiple rounds of 2D and 3D classification, a 3D refined cryo-EM density map with an overall resolution of 3.4 Å was obtained from 333,051 particles. To avoid the potential averaging out of asymmetric features due to C2 symmetry, C1 symmetry was adopted in the 3D refinement. For the analysis of OsPT11^pH8^, a similar process was followed, enriching 346,493 particles. After 3D refinement, a cryo-EM density map of OsPT11^pH8^ with an overall resolution of 3.8 Å was obtained. The FSC curve indicated an overfitting in refinements, suggesting that bad particles might have affected the 3D refinement. Therefore, the particles were re-extracted into a 280-pixel box, obtaining 345,782 particles, which were further subjected to 3D classification. One class containing 175,967 particles was refined, resulting in a cryo-EM density map with an overall resolution of 3.4 Å. The resolution of all the cryo-EM maps reported above are based on the gold-standard Fourier shell correlation (FSC)^56^. Local resolution estimation was performed using MonoRes^57^.

### Model building and refinement

The initial model was predicated from AlphaFold2^58^. We employed the ChimeraX (v1.6.1) software to dock this predicted model into the reconstructed cryo-EM density map. The model was manually refined through iterative rounds of adjustments using COOT^59^. The obtained model was refined against the map in real space using PHENIX with the application of secondary structure and geometric constraints^60^. Model quality was assessed using MolProbity scores^61^, Ramachandran plots, and EMRinger^62^. Figures were generated using ChimeraX (v1.6.1) and PyMol (v.2.4.1).

### Yeast Complementation

For yeast complementation, the coding regions of yeast PHO84, yeast PHO91, and OsPT11 were separately inserted into the PRS416-ADH vector and transformed into the yeast Pi transport-deficient mutant YP100 strain. The complemented genes were expressed under the control of the ADH promoter. Transformants were selected on synthetic complete (SC-uracil) medium containing 2% galactose, 0.67% yeast nitrogen base (YNB) without amino acids, 0.2% appropriate amino acids, and 2% agar, and incubated at 28 °C for 3-4 days. Yeast strains were verified by PCR, and then grown in liquid 2× YPD medium. Mid-exponential phase cells were collected, washed twice with water, and resuspended to an OD_600_ of 1. Equal volumes of five serial dilutions were spotted onto YNB medium (without phosphate) containing 2% glucose, 2% agar, appropriate amino acids, and varying Pi concentrations at pH 5.4. Plates were incubated at 28 °C for 4-5 days. For growth curve analysis, yeast cultures were resuspended to OD_600_ = 1, diluted 1:50 in liquid YNB, and incubated at 28°C for 48 hours in a FLUOstar OMEGA plate reader (BioDOT) for measurement.

### Electrophysiological experiments

To construct the oocyte expression vectors for electrophysiology studies, cDNAs of OsPT2, OsPT11, and point mutations were cloned into the pNB1-YFP expression vectors using the USER method^63^. Uracil-containing forward primer was designed as 5’-GGCTTAAU + sequence complementary to the target gene-3’, and reverse primer as 5’-GGTTTAAU + sequence complementary to the target gene-3’. PCR was performed with Phusion U Hot Start DNA polymerase (Thermo Fisher, F555) according to the manufacturer’s instructions. The reaction mixture containing PCR product, USER enzyme mix (New England Biolabs, M550), and PacI/Nt.BbvCI digested pNB1 vector was incubated for 20 minutes at 37 °C followed by 20 minutes at 25 °C, and then transformed into chemically competent *E. coli*.

Capped RNA (cRNA) was prepared from the linearized pNB1-YFP vector DNA templates using the mMESSAGE mMACHINE^®^ high yield capped RNA T7 kit following the manufacturer’s instructions (AmbionTM USA). The quality of cRNAs was checked by Nanodrop, and the concentration was adjusted to the same level and stored at -80 °C until injection. The protein expression and two-electrode voltage-clamp recordings in *Xenopus* oocytes were performed as described previously^64^. *Xenopus* oocytes were harvested at stage V to VI and maintained in ND96 solution (96 mM NaCl, 2 mM KCl, 1 mM CaCl_2_, 1 mM MgCl_2_, 5 mM HEPES, 10 mM sorbitol, pH was adjusted to 7.4 with NaOH) for overnight prior to cRNA-injections. Each oocyte was injected with 30 ng cRNAs or the same amount of water. Injected oocytes were incubated in ND96 solution at 18 °C for 1.5 days, transferred to ND96 solution with 5 mM KH_2_PO_4_, and incubated overnight. Protein expression was detected by YFP fluorescence signals (excited at 514 nm) using the Leica SP8 Laser confocal microscope (Leica^TM^ GER). The currents were recorded with hyperpolarized pulses of a 0.1-s pre-pulse at -40 mV, followed by voltage steps of +80 to -180 mV (step at -20 mV, 1.5 s duration) and a 0.5 s deactivation at -40 mV using Axon Axoclamp 900A Amplifier and Digidata 1440A low-noise data acquisition system controlled by Clampex10.4 acquisition software. The bath solution contained 0, 10, or 30 mM KH_2_PO_4_, 10 mM NaCl, 4 mM KCl, 1 mM CaCl_2_, 1 mM MgCl_2_, 10 mM HEPES, pH was adjusted to 5.5 or 7.5 with NaOH, and osmolality was adjusted to 220 mOsm/L with mannitol. The pipette solution contained 3 M KCl.

### Proteoliposome-based transport assay

For reconstitution of OsPT11 proteoliposomes, 10 mg/mL *E. coli* total extract (Avanti Polar Lipids) was solubilized in reconstitution buffer (10 mM HEPES-Tris, pH 7.4, and 100 mM KCl). Preformed liposomes were dissolved in 1.3% (w/v) DDM and mixed with purified OsPT11 or its variants at a protein-to-lipid ratio of 1:100 (w/w). After incubating at 4 °C for 1.5 hours, DDM was removed by adding SM-2 bio-beads (Bio-Rad) at a DDM-to-bio-bead ratio of 1:20 (w/w) and incubating overnight. The resulting proteoliposomes were extruded through a 200 nm polycarbonate filter (Whatman) for the transport assay.

For the transport assay, 15 μL of proteoliposome containing 0.2–0.5 μg of protein were diluted into 80 μL assay buffer (10 mM MES, pH 6.0, and 100 mM KCl). Pi transport reactions were initiated by adding a KPi mixture at given concentration, which contains non-labeled KH_2_PO_4_ and [^32^P] KH_2_PO_4_ (3.7 MBq/ μmol; PerkinElmer) in a molar ratio of 29:1-3:1. The assays were performed at 37 °C, and terminated at given times by diluting tenfold with ice-cold stop buffer (10 mM Hepes-Tris, pH 7.4, 100 mM KCl and 5 mM non-labeled KH_2_PO_4_), followed by rapid filtration through nitrocellulose membrane (Millipore, 0.22 μm Triton-free MCE). The filters were subsequently washed with 2×15 ml ice-cold stop buffer, placed in 5 mL Optiphase HiSafe 3 scintillation fluid and counted after 14 hours. Background counts were determined from parallel transport assays that were terminated at the start of the reaction. After subtracting the background, the amount of phosphate transported into the proteoliposomes was quantified by comparison to a standard curve of ^32^P KH_2_PO_4_. This standard curve was established by diluting sole ^32^P KH_2_PO_4_ in water and recording the corresponding radioactive counts. It thus provided a quantitative relationship between radioactive counts and the amount of ^32^P, which was used to quantify the amount of Pi taken up during our transport reactions. The protein content in the proteoliposomes was resolved by SDS-PAGE and quantified using ImageJ. The phosphate uptake was expressed as pmol Pi/μg protein. Each assay was independently performed a minimum of three times to generate an overall mean and standard error of the mean (SEM).

### Single-molecule fluorescence resonance energy transfer analysis

Maleimide-LD555 and maleimide-LD655 (Lumedyne Technologies) were used as the FRET donor and acceptor, respectively. These dyes were dissolved in DMSO to a final concentration of 1 mM prior to conjugation. Maleimide-LD555 and maleimide-LD655 were premixed and incubated with the protein sample at a molar ratio of 4:8:1. Conjugation was carried out for 30 minutes at room temperature in dark. Excess dyes were removed using a desalting column equilibrated with desalting buffer (25 mM Tris-HCl, pH 8.0, 100 mM NaCl, and 0.06% (w/v) digitonin), and the target protein was collected for single-molecule FRET data acquisition.

Microfluidic imaging chambers were passivated with a mixture of polyethylene glycol (PEG) and biotinylated PEG (Laysan Bio) and incubated for 2 minutes with 0.01 mg/mL streptavidin (Invitrogen), followed by 8 mg/mL BSA in buffer containing 150 mM NaCl and 25 mM Tris-HCl (pH 8.0). After a 10-minute incubation, the BSA was removed by washing the chambers with buffer A (25 mM Tris-HCl, pH 8.0, 150 mM NaCl, and 0.06% (w/v) digitonin). The dyes-labeled and biotinylated sample was then introduced at a final concentration of ∼0.1 nM and incubated for 2 minutes. Unbound protein was washed away with buffer A. Imaging was performed in deoxygenated buffer A, by including 0.8% (m/m) D-glucose, 26.6 U/mL glucose oxidase, 140 U/mL catalase, 5 mM protocatechuic acid, and 50 nM protocatechuate-3,4-dioxygenase, to minimize photobleaching.

Single-molecule experiments were conducted at room temperature using a custom-designed objective-type total internal reflection fluorescence microscope, based on a Nikon ECLIPSE Ti2 inverted microscope. This setup was equipped with a 100× oil objective (Nikon, 1.49 NA), an EMCCD camera (Andor iXon Ultra 897), and solid-state 532 nm excitation laser (Coherent Inc. OBIS Smart Lasers). Fluorescence emission from the probes was collected by the microscope and spectrally separated using an interference dichroic (T635lpxr, Chroma) in conjunction with bandpass filters (ET585/65m and ET690/70m, Chroma), all integrated within a Dual-View spectral splitter (Photometrics). The hardware was controlled, and single-molecule FRET (smFRET) movies were acquired using Cell Vision software (Beijing Coolight Technology).

Single-molecule fluorescence-time trajectories were extracted and filtered using the smCamera software with the following criteria: single-step donor photobleaching, total intensity ∼5000 Counts, <4 donor blinking events, and FRET efficiency above baseline for at least 100 frames. The extracted trajectories were subsequently analyzed using vbFRET^65^. The FRET efficiency was calculated as (IA-ID*0.07)/(0.98*ID+IA), where ID and IA represent the donor and acceptor fluorescence intensities, respectively. The crosstalk value of 0.07 was determined from a protein sample containing the donor fluorophore only, while the gamma factor value of 0.98 represents the ratio of the quantum yields and the detection efficiencies of donor and acceptor. The FRET efficiency histogram was calculated from > 100 fluorescence trajectories, fitted with a two-Gaussian peaks.

The accessible volume of conjugated fluorophores on the Pi-bound OsPT11 structure was sampled using the previously established method^40^. Briefly, to model the fluorescent probes on the Pi-bound OsPT11 structure, fluorophores were added to the labeling sites using Xplor-NIH (v3.6)^66^, and the linker between the protein backbone and the rigid portion of the fluorophore was given torsion angle freedom and were allowed reorient. A total of 456 structures were selected based on their overall energy, and the average distance between the geometric centers of the two chromophores was calculated, as shown in Supplementary Fig. 8.

## Data availability

The EM maps and atomic models of OsPT^pH5^ and OsPT^pH8^ have been deposited in the Electron Microscopy Data Bank and Protein Data Bank, with the accession numbers EMD-62480 and EMD-62441, and 9KOU.PDB and 9KMQ.PDB, respectively. Materials are available from the corresponding authors on request.

## Acknowledgments

We thank the Cryo-EM Center, the University of Science and Technology of China (USTC), for the EM facility support. We are grateful to Dr. Yongxiang Gao (USTC) for technical support during EM image acquisition. We thank the Cryo-EM Facility of Southern University of Science and Technology for providing technical support. We thank the Center for Protein Research, and Dr. Jianbo Cao at the Public Laboratory of Electron Microscopy, Huazhong Agricultural University, for technical support. This work was supported by the National Natural Science Foundation of China (32422041 to Z.L., 32071226 to Z.L., 32201005 to Z.G., 32070214 to S.X., and 31670267 to S.X.), the Foundation of Hubei Hongshan Laboratory (2021HSZD011 to P.Y.), the Fundamental Research Funds for the Central Universities (2662023PY001 to P.Y.) and the HZAU-AGIS Cooperation Fund (SZYJY2022022 to Z.L.). Z.G. acknowledges the support of National Postdoctoral Program for Innovative Talents (BX2021108). H.L. acknowledges the support of the Postdoctoral Fellowship Program of CPSF (GZC20230908) and the China Postdoctoral Science Foundation (CN: 2023M741289).

## Author contributions

Z.L., Y.C., and S.X. conceived and supervised the project. Z.D., Z.G., H.L., J.Z., and H.H. designed experiments. Z.D. prepared samples. Z.G. determined the structures. H.L. performed electrophysiological experiments. J.Z. and H.H. performed smFRET analysis. Z.D. and Y.C. performed proteoliposome-based transport assay. Z.Z., W.Z., L.H., J.Z., Y.L., B.W. and H.T. contributed to plasmids constructing and data collecting. F.D., X.L., L.X. and P.Y. contributed to the data analysis. Z.L. and Y.C. wrote the manuscript with help from all authors.

---

**Supplementary Fig. 1.**
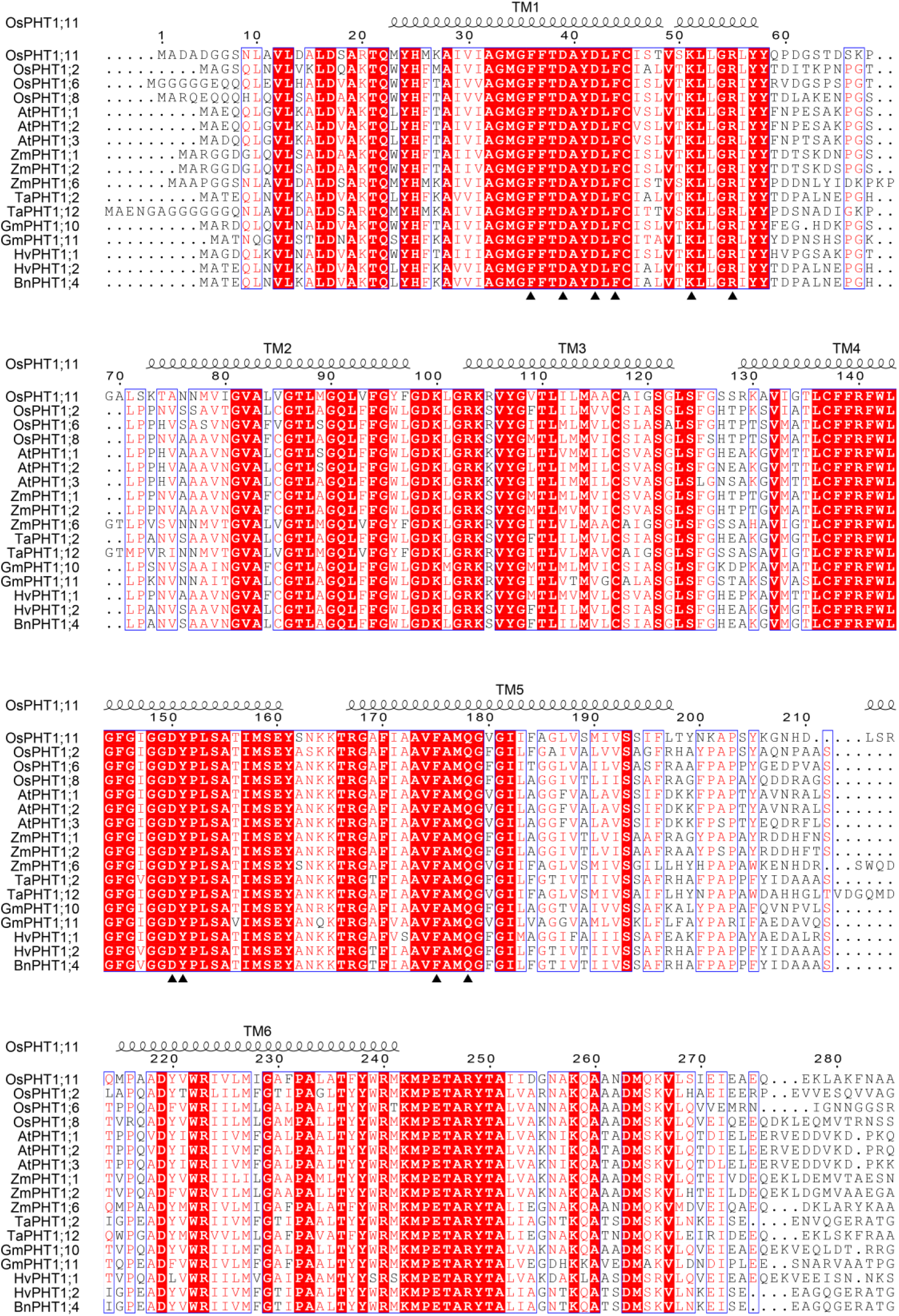
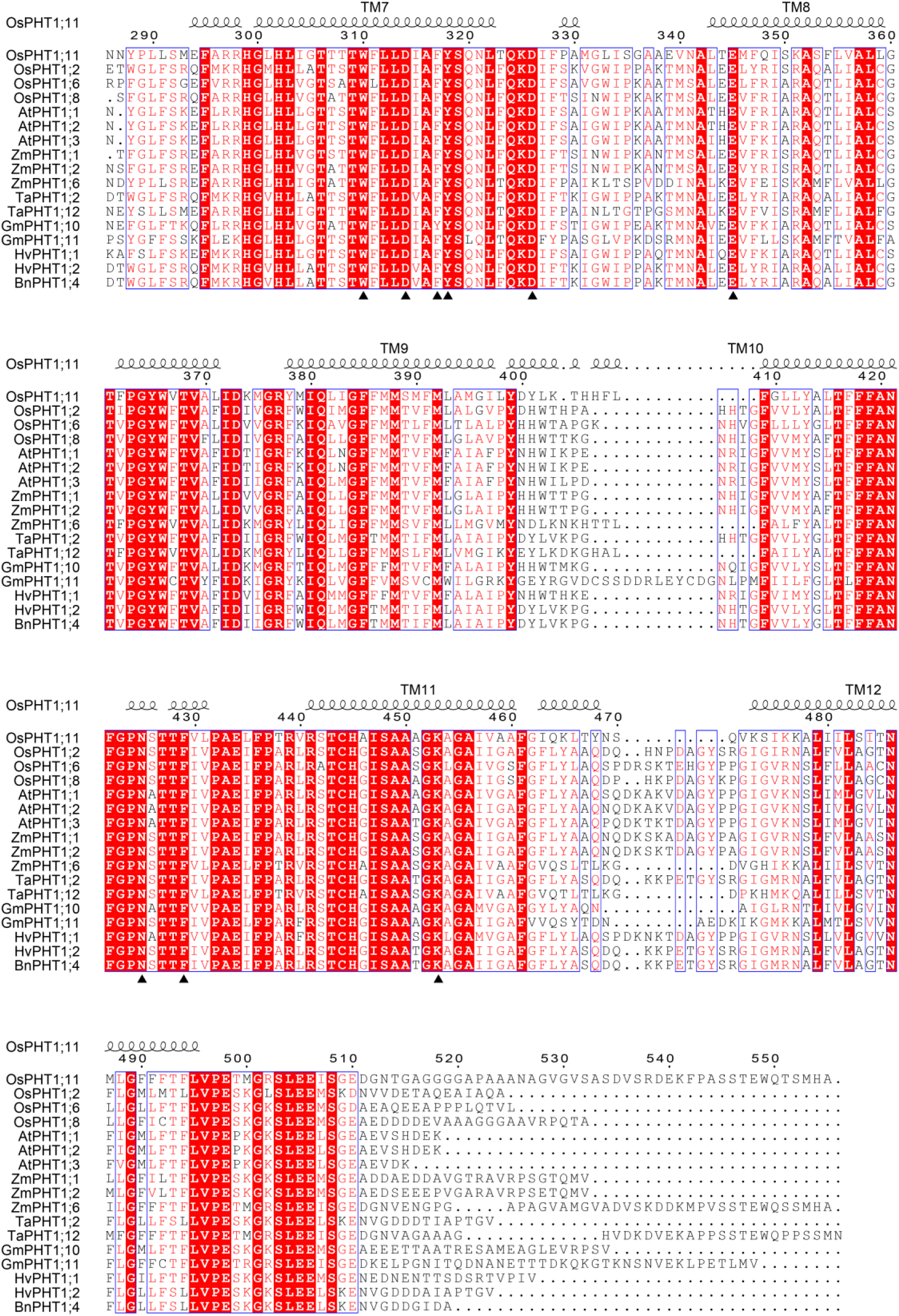
Sequence alignment of representative plant PHT1 transporters. The sequence of rice OsPHT1;11 (OsPT11) is aligned with other orthologues. Structure-based alignment was performed by ESPript (3.0). The sequence identity is indicated by white letters against a red background, and the sequence of a similarity over 90% is indicated by red letters. The secondary elements of OsPHT1;11 are labeled at the top of the alignment. The conserved residues mentioned in the main text are marked with black triangles at the bottom of the alignment.

**Supplementary Fig. 2.**
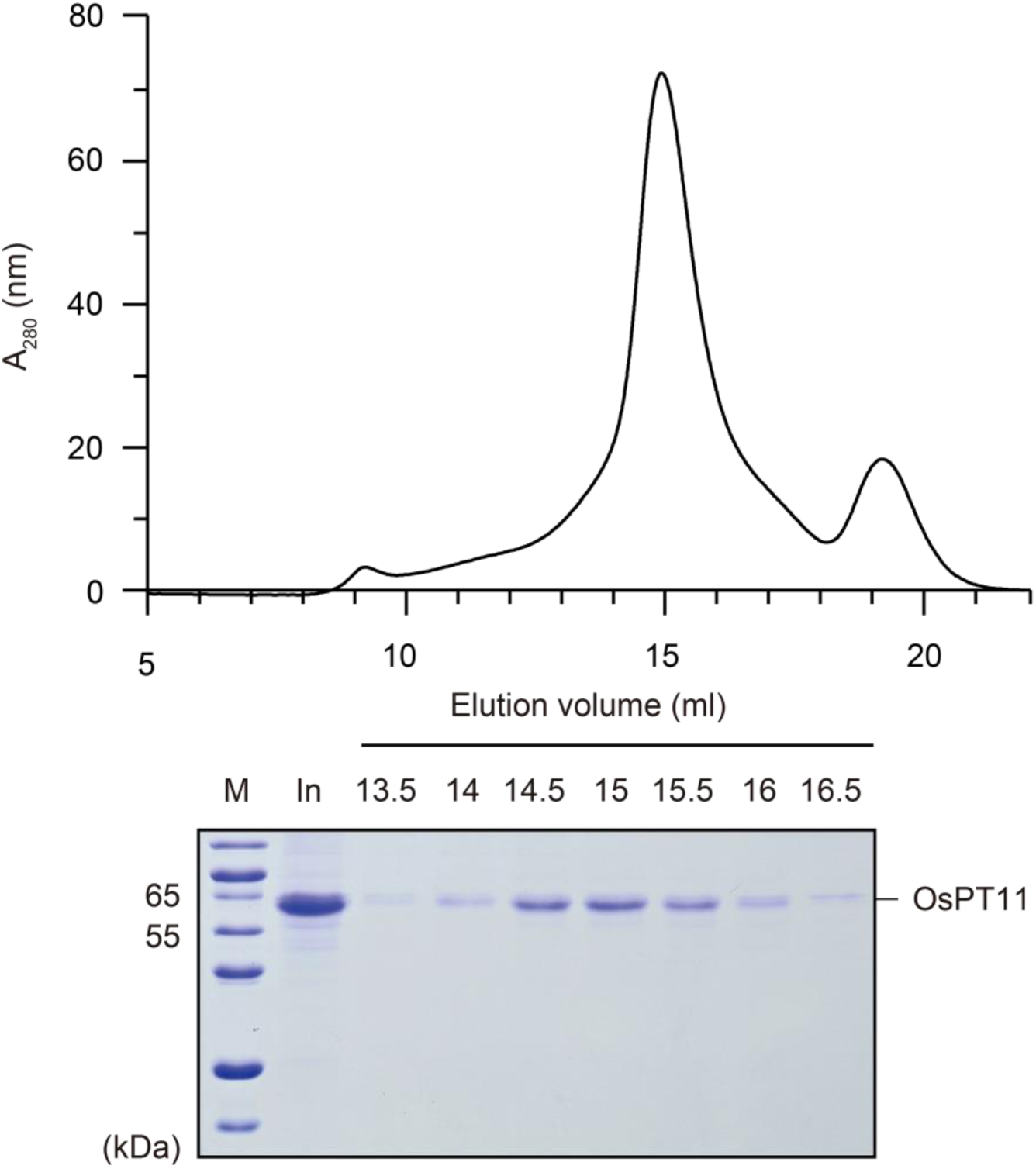
Representative size-exclusion chromatography (SEC) purification of OsPT11. Fractions were resolved by 15% SDS-PAGE, and the peak fractions were collected and concentrated for further use. In, input sample before SEC step.

**Supplementary Fig. 3.**
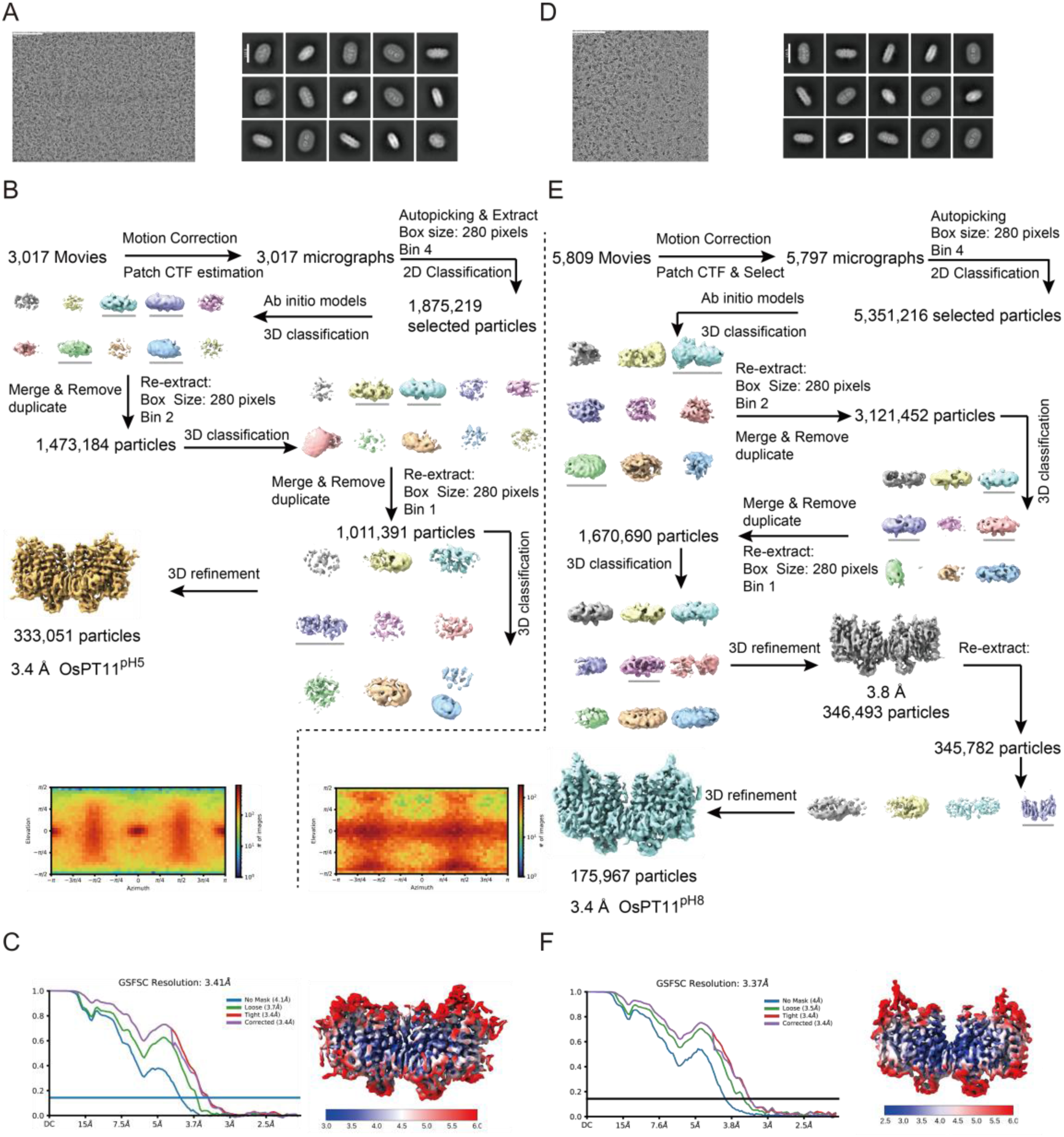
Cryo-EM data processing workflow. Cryo-EM data processing for OsPT11^pH5^ (**A-C**) and OsPT11^pH8^ (**D-F**). (**A, D**) Representative micrographs and 2D class averages. (**B, E**) Cryo-EM data processing workflow. (**C, F**) Gold standard Fourier shell correlation (FSC) curves (left panel) and local resolution (right panel) from 3D refinement for the final reconstructions.

**Supplementary Fig. 4.**
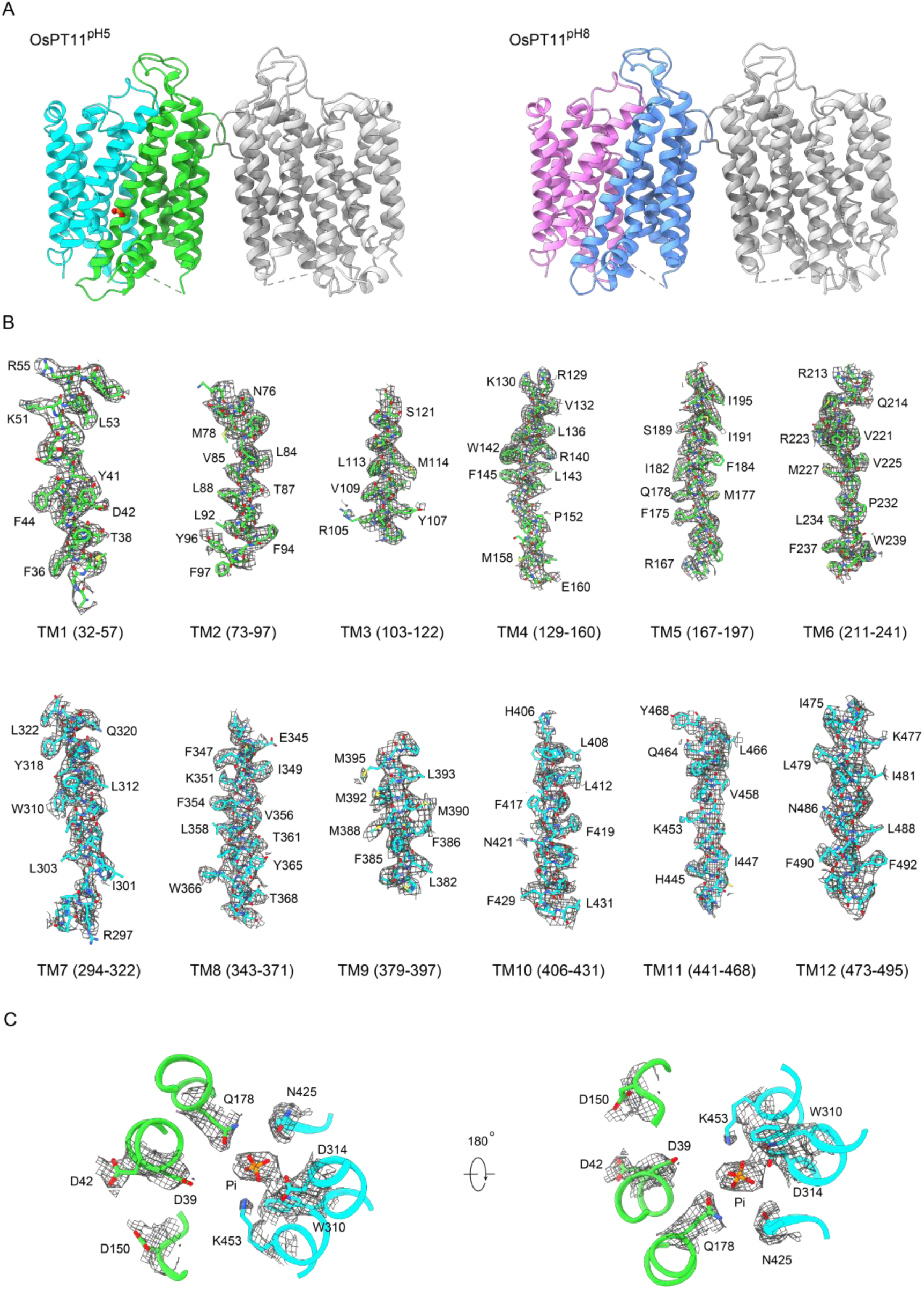
Structures of OsPT11 and EM densities of representative elements. **A** Cartoon representation of the resolved dimeric structure of OsPT11 determined at pH 5 and pH 8, respectively. The phosphate bound to one protomer of OsPT11 at pH 5 is represented as a sphere. **B** EM density maps for the indicated transmembrane α-helices of Pi-bound protomer in OsPT11^pH5^ structure. Map contour level = 0.35-0.40 in ChimeraX. **C** EM density maps of the Pi binding site in the Pi-bound protomer in OsPT11^pH5^ structure. This figure corresponds to Fig. 3A in the main text. Map contour level = 0.34 in ChimeraX.

**Supplementary Fig. 5.**
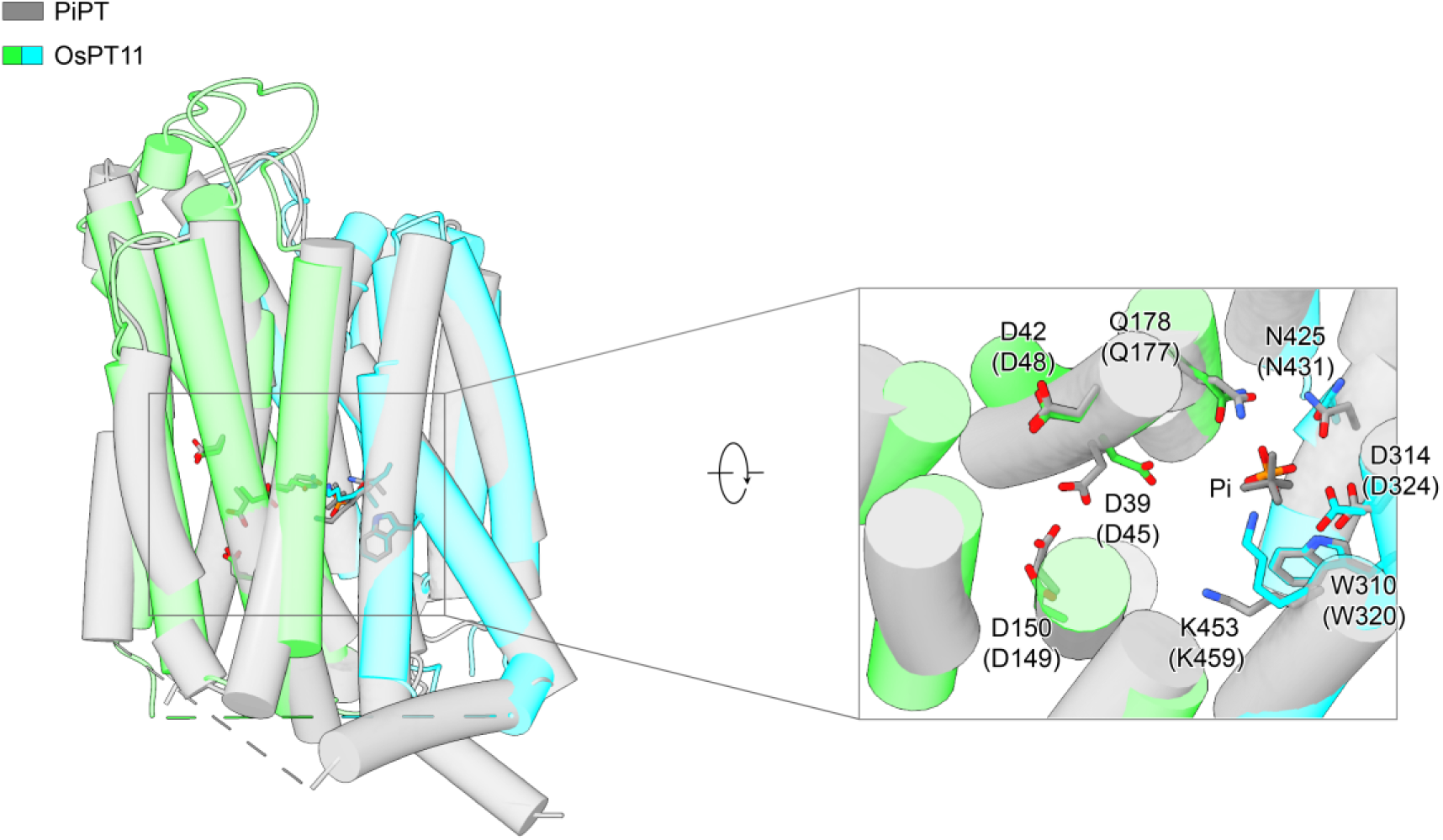
Structural comparison between Pi-bound PiPT (7SP5.PDB) and OsPT11. Corresponding residues in PiPT are indicated in brackets. The two structures share similarities, with an overall RMSD of 1.1 Å, suggesting that the Pi-bound occluded conformational state is structurally conserved across fungi and higher plants.

**Supplementary Fig. 6.**
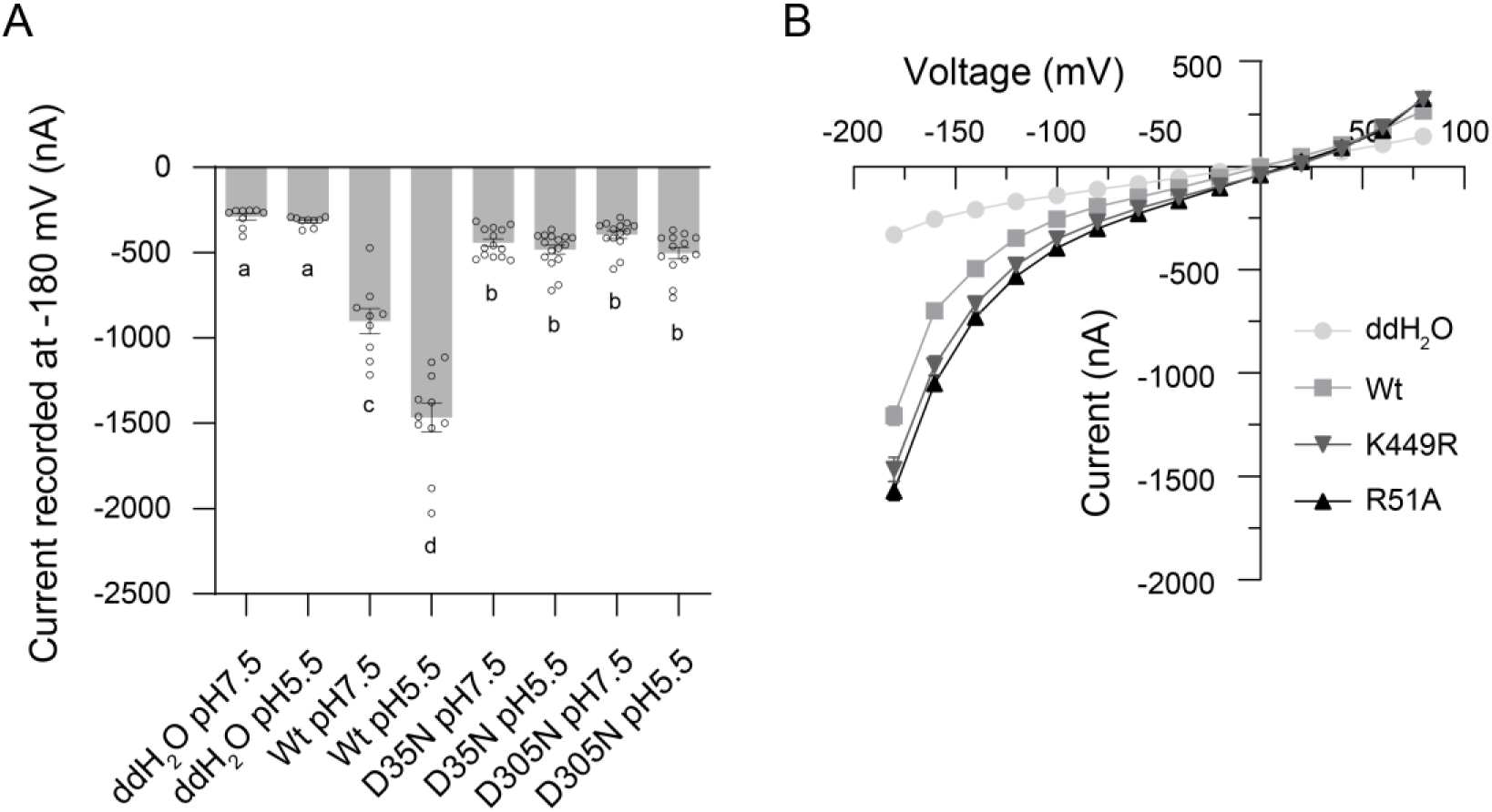
Electrophysiological experiments of OsPT2 and variants. **A** Currents were recorded at −180 mV in *Xenopus* oocytes injected with water (n=6, pH 7.5; n=6, pH 5.5), expressing OsPT2 (Wt, n=9, pH 7.5; n=11, pH 5.5), D35N mutant (n=14, pH 7.5; n=16, pH 5.5), D305N mutant (n=14, pH 7.5; n=13, pH 5.5) in the bath solution with 30 mM Pi. Data are represented as mean ± SEM. Different letters represent p < 0.05; One-way ANOVA. **B** Current-voltage relationship was recorded in oocytes injected water (n=6), expressing OsPT2 (Wt, n=16), R51A (n=18), K453R (n=24) in the presence of 30 mM Pi at pH 5.5. Data are represented as mean ± SEM.

**Supplementary Fig. 7.**
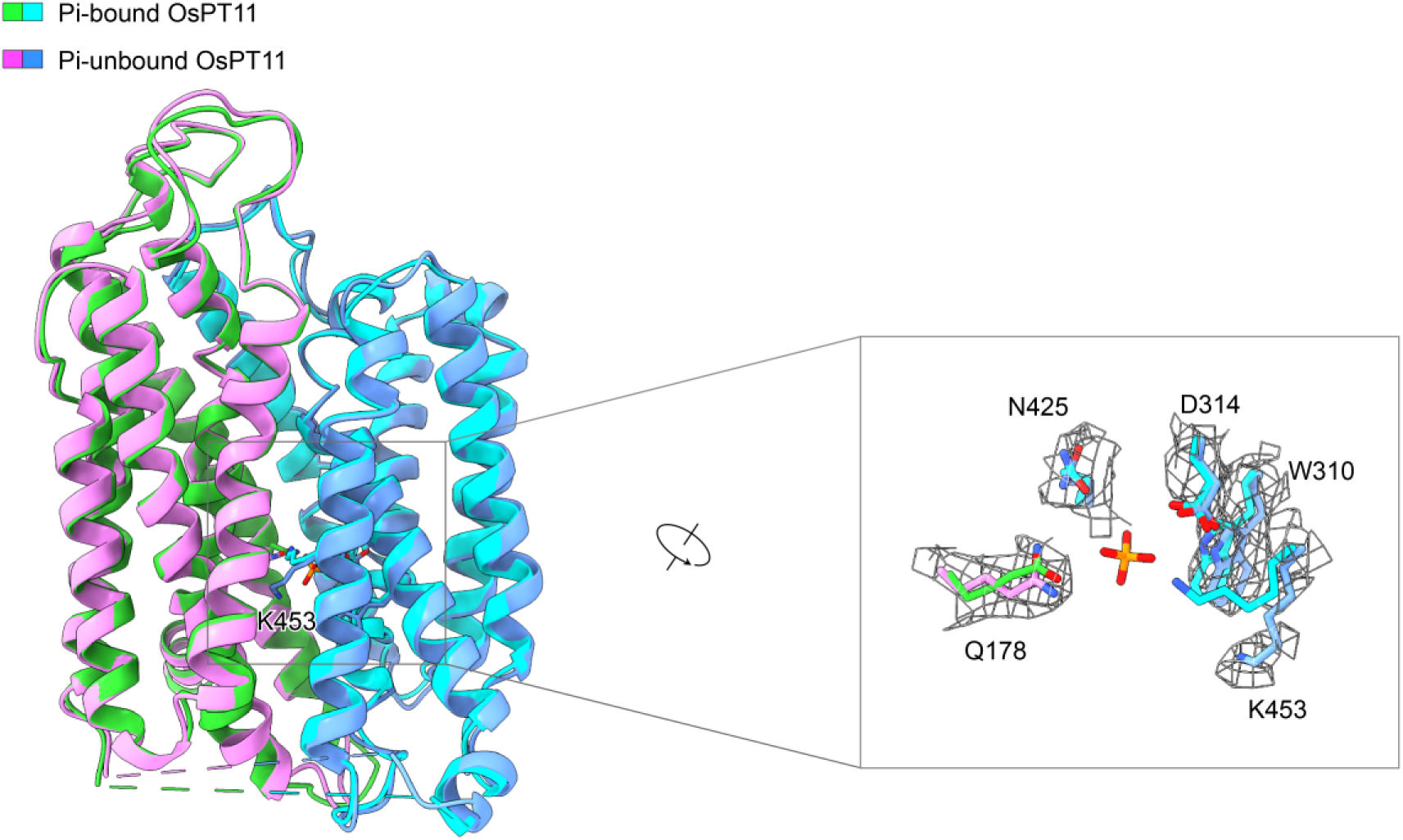
Structural comparison between the Pi-bound and unbound forms of OsPT11. The overall RMSD of the two structures is 0.7 Å. EM density maps of the Pi binding site in the Pi-unbound OsPT11 structure are shown with a contour level of 0.32 in ChimeraX. EM density maps of the Pi binding site in the Pi-bound OsPT11 structure can be found in Supplementary Fig. 4C.

**Supplementary Fig. 8.**
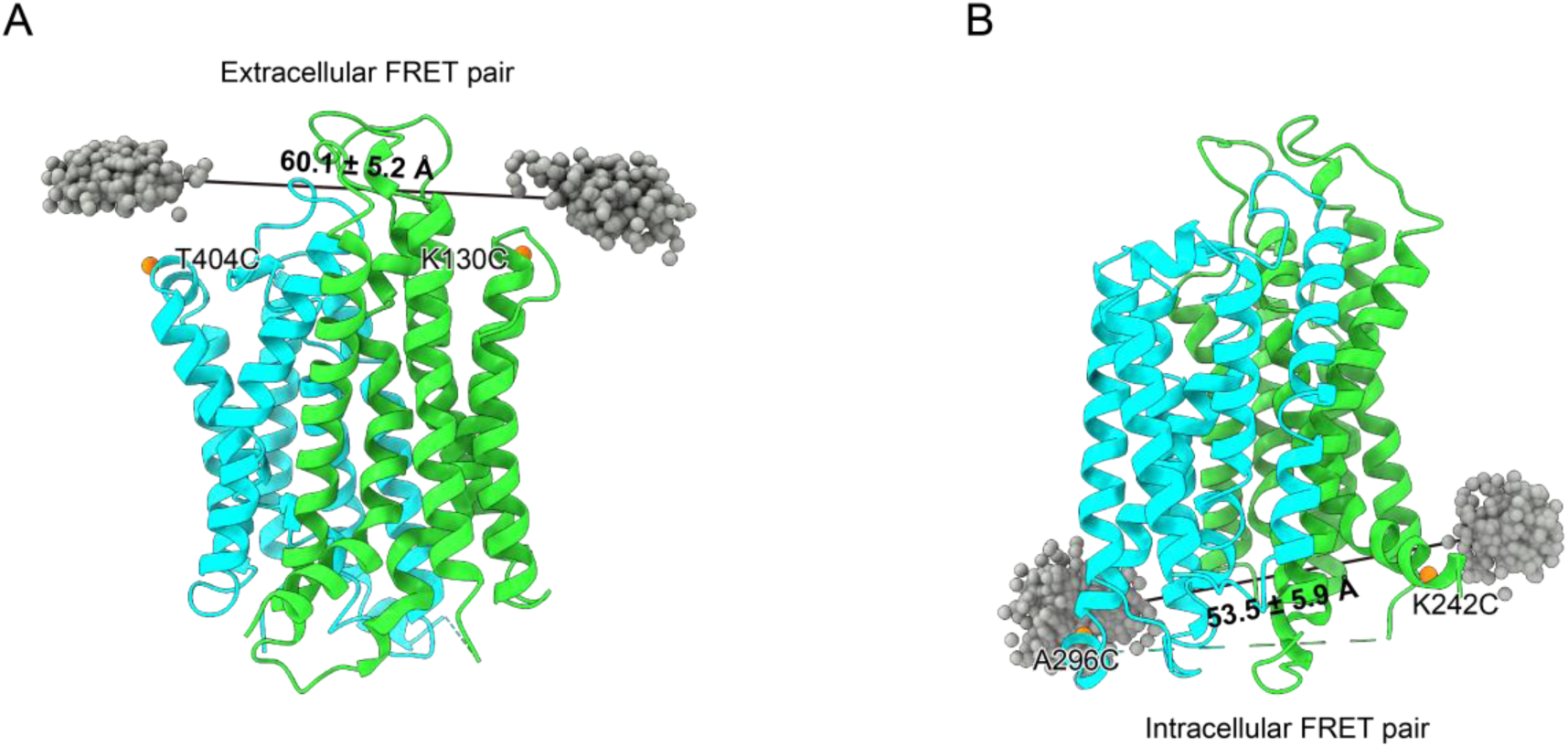
Calculated average distance between conjugated probes. The accessible volume of conjugated fluorophores on the occluded OsPT11 structure were sampled using Xplor-NIH (See Methods for details). The geometric centers of the two chromophores were represented with gray spheres, and the average distance between them is 60.1 ± 5.2 Å for the extracellular FRET pair (**A**), and 53.3 ± 5.9 Å for the intracellular FRET pair (**B**).

**Supplementary Table 1.** Cryo-EM data collection, refinement and validation statistics.

|  | OsPT11<sup>pH5</sup><br>(EMD-62480)<br>(PDB 9KOU) | OsPT11<sup>pH8</sup><br>(EMD-62441)<br>(PDB 9KMQ) |
| --- | --- | --- |
| **Data collection and processing** |  |  |
| Magnification | 81,000 | 130,000 |
| Voltage (kV) | 300 | 300 |
| Camera | K3 | K2 |
| Electron exposure (e<sup>-</sup>/Å<sup>2</sup>) | 50 | 50 |
| Defocus range (μm) | -1.2~-2.2 | -1.2~-2.2 |
| Pixel size (Å) | 1.07 | 1.08 |
| Micrographs collected (no.) | 3,017 | 5,809 |
| Micrographs used (no.) | 3,017 | 5,797 |
| Total extracted particles (no.) | 2,305,826 | 5,454,082 |
| Symmetry imposed | C1 | C1 |
| Refined particle images (no.) | 333,051 | 175,967 |
| Map resolution (Å) | 3.4 | 3.4 |
| FSC threshold | 0.143 | 0.143 |
| Map resolution range (Å) | 3-6.5 | 2.50-6.05 |
| **Refinement** |  |  |
| Initial model used | AlphaFold2 | AlphaFold2 |
| Model resolution (Å) | 3.38/3.71 | 3.3/3.59 |
| FSC threshold | 0.143/0.5 | 0.143/0.5 |
| Map sharpening *B* factor (Å<sup>2</sup>) | 148.7 | 127.3 |
| Model composition |  |  |
| Non-hydrogen atoms | 6,672 | 6,676 |
| Protein residues | 861 | 862 |
| Ligands | 1 | 0 |
| *B* factors (Å<sup>2</sup>) |  |  |
| Protein | 90.28 | 78.37 |
| Ligand | 85.88 | NA |
| R.m.s. deviations |  |  |
| Bond lengths (Å) | 0.004 | 0.003 |
| Bond angles (°) | 0.835 | 0.683 |
| Validation |  |  |
| MolProbity score | 2.05 | 2.39 |
| Clashscore | 11.80 | 9.64 |
| Poor rotamers (%) | 0.87 | 4.19 |
| Ramachandran plot |  |  |
| Favored (%) | 92.61 | 93.91 |
| Allowed (%) | 7.39 | 6.09 |

| Disallowed (%) | 0 | 0 |
| --- | --- | --- |

